# Genetic structure and molecular diversity of Brazilian grapevine germplasm: management and use in breeding programs

**DOI:** 10.1101/2020.05.05.078865

**Authors:** Geovani Luciano de Oliveira, Anete Pereira de Souza, Fernanda Ancelmo de Oliveira, Maria Imaculada Zucchi, Lívia Moura de Souza, Mara Fernandes Moura

**Affiliations:** Advanced Fruit Research Center, Agronomic Institute (IAC), Jundiaí, SP, Brazil; Molecular Biology and Genetic Engineering Center (CBMEG), University of Campinas (UNICAMP), Campinas, SP, Brazil; Department of Plant Biology, Biology Institute, University of Campinas (UNICAMP) UNICAMP, Campinas, SP, Brazil; Laboratory of Conservation Genetics and Genomics, Agribusiness Technological Development of São Paulo (APTA), Piracicaba, SP, Brazil

## Abstract

The management of germplasm banks is complex, especially when many accessions are involved. Microsatellite markers are an efficient tool for assessing the genetic diversity of germplasm collections, optimizing their use in breeding programs. This study genetically characterizes a large collection of 410 grapevine accessions maintained at the Agronomic Institute of Campinas (IAC) (Brazil). The accessions were genotyped with 17 highly polymorphic microsatellite markers. Genetic data were analyzed to determine the genetic structure of the germplasm, quantify its allelic diversity, suggest the composition of a core collection, and discover cases of synonymy, duplication, and misnaming. A total of 304 alleles were obtained, and 334 unique genotypes were identified. The molecular profiles of 145 accessions were confirmed according to the literature and databases, and the molecular profiles of more than 100 genotypes were reported for the first time. The analysis of the genetic structure revealed different levels of stratification. The primary division was between accessions related to *Vitis vinifera* and *V. labrusca*, followed by their separation from wild grapevine. A core collection of 120 genotypes captured 100% of all detected alleles. The accessions selected for the core collection may be used in future phenotyping efforts, in genome association studies, and for conservation purposes. Genetic divergence among accessions has practical applications in grape breeding programs, as the choice of relatively divergent parents will maximize the frequency of progeny with superior characteristics. Together, our results can enhance the management of grapevine germplasm and guide the efficient exploitation of genetic diversity to facilitate the development of new grape cultivars for fresh fruits, wine, and rootstock.

## Introduction

Grapevine (*Vitis* spp.) is considered to be a major fruit crop globally based on hectares cultivated and economic value [1]. Grapevines are exotic species in Brazil but have become increasingly important in national fruit agriculture in recent years, transitioning from exclusive cultivation in temperate zones to a great alternative in tropical regions.

European grapevine, or *V. vinifera*, cultivars stand out in terms of their economic importance, being the most commonly planted worldwide and characterized by having fruits of excellent quality with wide morphological and genetic diversity. They are widely used for the production of fresh fruits, dried fruits, and juice and in the global fine and sparkling wine industry [2].

In Brazil, the American *V. labrusca* varieties and hybrids (*V. labrusca* x *V. vinifera*) thrive because of their vegetative characteristics, which are best adapted to the country’s environmental conditions, with generally high humidity. In addition, due to their relatively high robustness, they are resistant to many diseases that affect grapevine in the country, resulting in production of relatively high volume, although of low quality, and have become dominant on Brazilian plantations [3, 4].

The wild species of the genus *Vitis* have contributed evolutionarily through interspecific crossings, accidental or planned, to the adaptation of grapevine to the highly different conditions that its expansion has demanded. Hybrid varieties are characterized by greater resistance to pests and diseases than *V. vinifera* and by producing fruits with better organoleptic characteristics than American grapes. Crosses and natural mutations have greatly benefited from the possibility of vegetative propagation among grapevines, enabling the exploitation of different characteristics over time, with noticeable variations in berries, flowers, and leaves, further increasing the number of cultivars planted [2, 5].

The starting point of any breeding program of a species is genetic variability, whether spontaneous or created. The manipulation of this variability with suitable methods leads to the safe obtainment of superior genotypes in relation to agronomic characteristics of interest [6]. Germplasm banks have a fundamental role in preserving this genetic variability but require the maintenance of accessions [7]. The quantification of the magnitude of genetic variability and its distribution between and within the groups of accessions that constitute germplasm banks is essential to promote its rational use and adequate management [8].

Most germplasm is derived from seeds, but for highly heterozygous plants, such as grapevines, this method is not suitable, with conservation most commonly occurring through the use of *ex situ* field collections. The germplasm banks involved in breeding programs are fundamental to the development of new materials. These collections generally have a large number of accessions, but only a small proportion of these resources are used in practice. The management of such collections becomes complex when many accessions are involved. Redundancy should be reduced to a minimum, the use of “true-to-type” plant material must be ensured, and the introduction of new accessions should be optimized [9]. Therefore, it is essential to identify and correct errors related to synonyms, homonyms, and mislabeling that can occur during the introduction and propagation of plant material [10, 11]. The genetic characterization of available genetic resources may permit the optimization of the use of these resources by grouping a sufficient number of accessions in a core collection to maximize the genetic diversity described in the whole collection [12].

Information on the genetic diversity available in germplasm banks is valuable for use in breeding programs because such information assists in the detection of combinations of accessions capable of producing progenies with maximum variability in characteristics of interest, guiding hybridization schemes [13].

The identification of grapevine cultivars has traditionally been based on ampelography, which is the analysis and comparison of the morphological characteristics of leaves, branches, shoots, bunches, and berries [14], but as this process is carried out on adult plants, a long period is necessary before accession identification can be completed. Since many synonyms or homonyms exist for cultivars [2], passport data are not always sufficient to certify identities, mainly in terms of the distinction of closely related cultivars, and errors can arise. Thus, the use of molecular markers has become an effective strategy for this purpose due to the high information content detected directly at the DNA level without environmental influence and in the early stages of plant development, allowing for faster and more accurate cultivar identification [15].

Microsatellites, or simple sequence repeats (SSRs), are among the most appropriate and efficient markers for genetic structure and conservation studies [16]. SSRs are highly polymorphic and transferable among several species of the genus *Vitis* [17]. Since SSRs provide unique fingerprints for cultivar identification [18], they have been used for genetic resource characterization [19, 20], parentage analysis [21, 22], genetic mapping [23, 24], detection of quantitative trait loci (QTLs) [25], and assisted selection [26].

Because SSRs are highly reproducible and stable, they have allowed the development of several reference banks with grapevine variety genetic profiles from around the world. Access to these reference banks allows the exchange of information between different research groups, significantly increasing international efforts related to the correct identification of grapevine genetic resources [27].

Considering the importance of viticulture and winemaking in Brazil, the Agronomic Institute of Campinas (IAC) has a *Vitis* spp. germplasm bank including wild *Vitis* species, interspecific hybrids, and varieties of the main cultivated species (*V. vinifera*, *V*. *labrusca*, *V. bourquina*, *V*. *rotundifolia*) and varieties developed by the IAC. Our objective in the present study was to describe the diversity and genetic structure of the *Vitis* spp. available in this germplasm bank using microsatellite markers. The accessions were characterized, and their molecular profiles were compared with the use of different literature and online databases. Here we quantify the genetic diversity of this Brazilian germplasm and describe its genetic structure, and we suggest the composition of a core collection that would capture the maximum genetic diversity with a minimal sample size. We discuss perspectives related to the use of this information in germplasm management and conservation.

## Materials and Methods

### Plant material

A total of 410 accessions from the *Vitis* spp. Germplasm Bank of the IAC in Jundiaí, São Paulo (SP), Brazil, were analyzed. This germplasm encompasses more than ten species of *Vitis*, including commercial and noncommercial varieties of wine, table, and rootstock grapes. Each accession consisted of three clonally propagated plants, sustained in an espalier system and pruned in August every year, leaving one or two buds per branch. For sampling, were collected young leaves of a single plant from each accession. Detailed data on the accessions are available in S1 Table.

### DNA extraction

Total genomic DNA was extracted from young leaves homogenized in a TissueLyser (Qiagen, Valencia, CA, USA) following the cetyltrimethylammonium bromide (CTAB) method previously described by Doyle (1991) [28]. The quality and concentration of the extracted DNA were assessed using 1% agarose gel electrophoresis with comparison to known quantities of standard λ phage DNA (Invitrogen, Carlsbad, CA, USA).

### Microsatellite analysis

A set of 17 grapevine SSR markers well characterized in previous studies [22,29–31] were used, including ten developed by Merdinoglu et al. (2005) [32] (VVIn74, VVIr09, VVIp25b, VVIn56, VVIn52, VVIq57, VVIp31, VVIp77, VVIv36, VVIr21) and seven suggested by the guidelines of the European scientific community for universal grapevine identification, characterization, standardization, and exchange of information [33, 34]: VVS2 [35], VVMD5, VVMD7 [36], VVMD25, VVMD27 [37], VrZAG62, and VrZAG79 [38]. One primer in each primer pair was 5’ labeled with one of the following fluorescent dyes: 6-FAM, PET, NED, or VIC. Additional information about the loci is available in S2 Table.

Polymerase chain reaction (PCR) was performed using a three-primer labeling system [39] in a final volume of 10 μl containing 20 ng of template DNA, 0.2 μM of each primer, 0.2 mM of each dNTP, 2 mM MgCl2, 1× PCR buffer (20 mM Tris HCl [pH 8.4] and 50 mM KCl), and 1 U of Taq DNA polymerase. PCR amplifications were carried out using the following steps: 5 min of initial denaturation at 95°C followed by 35 cycles of 45 s at 94°C, 45 s at 56°C or 50°C (VVS2, VVMD7, VrZAG62 and VrZAG79), 1 min 30 s at 72°C, and a final extension step of 7 min at 72°C.

Amplifications were checked with 3% agarose gels stained with ethidium bromide. The amplicons were denatured with formamide and analyzed with an ABI 3500 (Applied Biosystems, Foster City, CA, USA) automated sequencer. The alleles were scored against the internal GeneScan-600 (LIZ) Size Standard Kit (Applied Biosystems, Foster City, CA, USA) using Geneious software v. 8.1.9 [40].

### Genetic diversity analyses

Descriptive statistics for the genotyping data were generated using GenAlEx v. 6.5 [41] to indicate the number of alleles per locus (Na), effective number of alleles (Ne), observed heterozygosity (H_O_), expected heterozygosity (H_E_), and fixation index (F). GenAlEx software was also used to identify private (Pa) and rare alleles (frequency < 0.05).

The polymorphism information content (PIC), discriminating power (D*j*), and null allele frequency (*r*) were calculated to evaluate the efficiency and discriminatory potential of each microsatellite marker. Polymorphism information content (PIC) was calculated using Cervus 3.0.7 [42] according to the expression 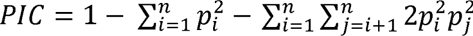, where *n* is the number of alleles, and *p_i_* and *p_j_* are the frequencies of the *i*^th^ and *j*^th^ alleles [43]. Discriminating power (D*j*) values were estimated to compare the efficiencies of microsatellite markers in varietal identification and differentiation. This parameter was calculated in accordance with the formula as follows: 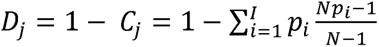, where D*_j_* is the probability that two randomly selected samples have different and distinct banding patterns, *p_i_* is the frequency of the *i*^th^ pattern revealed by each marker, N is the number of samples analyzed, and *I* is the total number of patterns generated by each marker [44].

The null allele frequency (*r*) was estimated using Cervus 3.0.7. By definition, a microsatellite null allele is any allele at a microsatellite locus that consistently fails to amplify to detectable levels via the polymerase chain reaction (PCR) [45]. Cervus 3.0.7 uses a iterative likelihood approach [46], in which the presence of null allele homozygotes is not taken into consideration initially but is added in later optimization rounds. This method avoids overestimating the frequency of a null allele if samples fail to amplify for reasons other than the presence of nulls [45].

### Genetic structure analysis

To assess the overall germplasm structuring, three approaches with different grouping criteria that do not require *a priori* assignment of individuals to groups were used: a Bayesian model-based approach, a distance-based model using a dissimilarity matrix, and discriminant analysis of principal components (DAPC).

The model-based Bayesian analysis implemented in the software package STRUCTURE v. 2.3.4 [47] was used to determine the approximate number of genetic clusters (K) within the full dataset and to assign individuals to the most appropriate cluster. STRUCTURE can identify subsets of individuals by detecting allele frequency differences within the data by assigning individuals to sub-populations based on analysis of likelihoods. The process begins by randomly assigning individuals to a pre-determined number of groups, after which variant frequencies are estimated in each group and individuals re-assigned based on those frequency estimates. This process is repeated many times in the burn-in process that results in a progressive convergence toward reliable allele frequency estimates in each population and membership probabilities of individuals to a population. During each analysis, membership coefficients summing to one are assigned to individuals for each group. If admixture is considered, membership coefficients are generated across multiple clusters. The assumptions are that loci are unlinked and populations are in Hardy-Weinberg Equilibrium (HWE) [48]. Additionally, a “hierarchical STRUCTURE analysis” [49] was applied in this study by running STRUCTURE subsequently for each identified cluster separately to reveal any underlying structure, as suggested by Pritchard et al. (2007) [50].

All simulations were performed using the admixture model, with 100,000 replicates for burn-in and 1,000,000 replicates for Markov chain Monte Carlo (MCMC) processes in ten independent runs. The number of clusters (K) tested ranged from 1 to 10.

The online tool Structure Harvester [51] was used to analyze the STRUCTURE output, and the optimal K values were calculated using Evanno’s ΔK *ad hoc* statistics [52]. The optimal alignment over the 10 runs for the optimal K values was obtained using the greedy algorithm in CLUMPP v.1.1.2 [53], and the results were visualized using DISTRUCT software v.1.1 [54]. Based on the posterior probability of membership (q), we classified individuals who showed q ≥ 0.70 as members of a given cluster. In contrast, accessions with a membership of q < 0.70 were classified as admixed. This procedure was performed to avoid individuals constrained to belong to any of the given number (K) of clusters.

Distance-based methods proceed by calculating a pairwise distance matrix, the entries of which provide the distance between every pair of individuals. This matrix may then be represented using some convenient graphical representation, such as a dendrogram, and clusters may be identified by eye [47]. Genetic distances between accessions were estimated on the basis of Rogers’ genetic distance [55], and the resulting distance matrix was used to construct a dendrogram with the neighbor-joining algorithm [56], with 1,000 bootstrap replicates implemented in the R package *poppr* [57]. The principle of this method is to find pairs of operational taxonomic units that minimize the total branch length at each stage of clustering starting with a star-like tree [56]. The final dendrogram was formatted with iTOL v. 5.5 [58].

DAPC as implemented in the R package *adegenet* 2.1.2 [59, 60] was also performed. DAPC is a multivariate analysis that does not rely on the assumption of HWE, the absence of linkage disequilibrium, or specific models of molecular evolution to identify clusters within genetic data. In DAPC, data are first transformed using a principal components analysis (PCA), after which a discriminant analysis (DA) is performed for the retained principal components. This process ensures that variables submitted to DA are perfectly uncorrelated and that their number is less than that of the analyzed individuals [61]. The *find.clusters* function was used to detect the number of clusters in the germplasm, which runs successive K-means clustering with increasing numbers of clusters (K). We used 20 as the maximum number of clusters. The optimal number of clusters was estimated using the Bayesian information criterion (BIC), which reaches a minimum value when the best-supported assignment of individuals to the appropriate number of clusters is approached. DAPC results are presented as multidimensional scaling plots.

### Accession name validation

To verify the trueness to type and identify misnamed genotypes, the molecular profiles obtained in this study were compared with the data contained in the following online databases: Vitis International Variety Catalogue (VIVC, www.vivc.de), Italian Vitis Database (http://www.vitisdb.it), “Pl@ntGrape, le catalogue des vignes cultivées en France” (http://plantgrape.plantnet-project.org/fr) and the U.S. National Plant Germplasm System (NPGS, https://npgsweb.ars-grin.gov/gringlobal/search.aspx). For this comparison, the molecular profile of seven microsatellite loci (VVS2, VVMD5, VVMD7, VVMD25, VVMD27, VrZAG62, VrZAG79) adopted by the databases was used.

The allele sizes were first standardized for consistency with various references [62]. If an accession was not listed in these databases, it was verified in other scientific papers.

### Core collection sampling

The R package *corehunter* 3.0 [63] was used to generate the core collection to represent the maximum germplasm genetic variability in a reduced number of accessions. Different samples were generated by changing the *size* parameter of the desired core collections to identify the subset of genotypes that could capture the entire diversity of alleles. The sizes ranged from 0.1 to 0.3 for all datasets. For each sample, the genetic diversity parameters were determined with GenAlEx v. 6.5 [41].

### Ethics statement

We confirm that no specific permits were required to collect the leaves used in this study. This work was a collaborative study performed by researchers from the IAC (SP, Brazil), São Paulo’s Agency for Agribusiness Technology (APTA, SP, Brazil), and the State University of Campinas (UNICAMP, SP, Brazil). Additionally, we confirm that this study did not involve endangered or protected species.

## Results

### Genetic diversity

Four hundred and ten grapevine accessions of *Vitis* spp. were analyzed at 17 SSR loci (S1 Table), and a total of 304 alleles were detected (Table 1). The number of alleles per SSR locus (Na) ranged from 10 (VVIq57) to 24 (VVIp31), with an average of 17.88. The number of effective alleles per locus (Ne) varied from 2.39 (VVIq57) to 11.40 (VVIp31), with a mean value of 7.02.

**Table 1.**
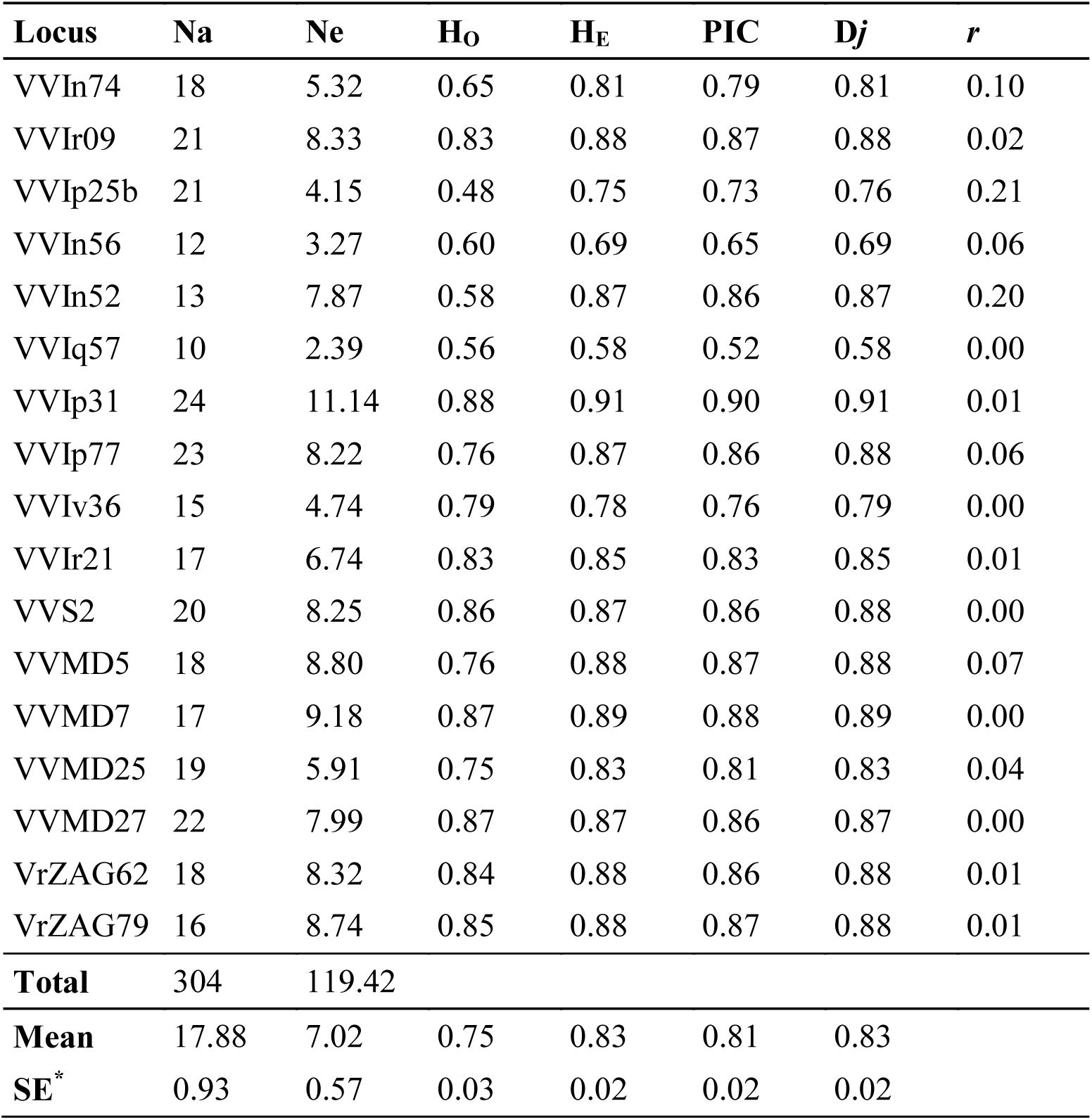
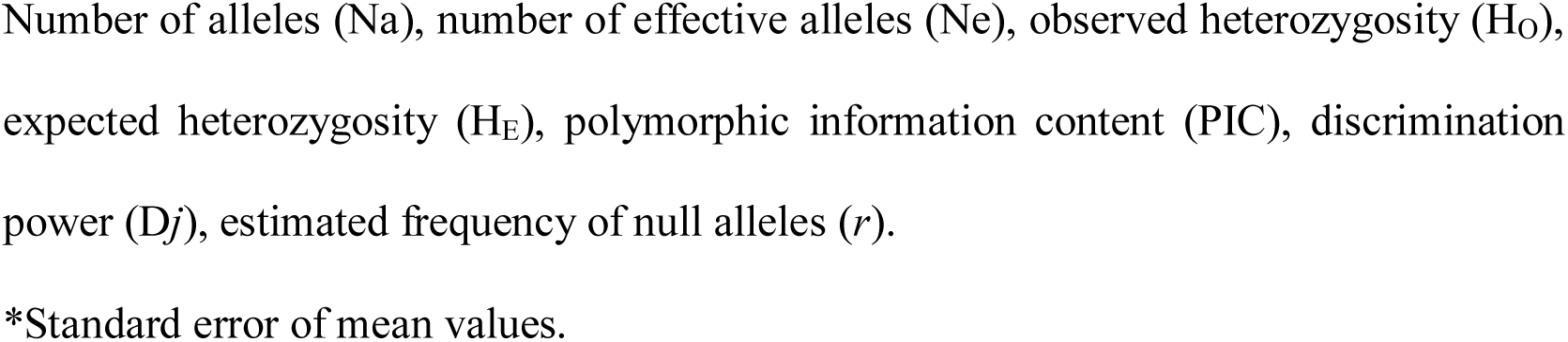
Genetic parameters of the 17 microsatellite loci obtained from 410 grapevine accessions.

| Locus | $N_a$ | $N_e$ | $H_o$ | $H_E$ | PIC | $D_j$ | $r$ |
| --- | --- | --- | --- | --- | --- | --- | --- |
| VVIn74 | 18 | 5.32 | 0.65 | 0.81 | 0.79 | 0.81 | 0.10 |
| VVIr09 | 21 | 8.33 | 0.83 | 0.88 | 0.87 | 0.88 | 0.02 |
| VVIp25b | 21 | 4.15 | 0.48 | 0.75 | 0.73 | 0.76 | 0.21 |
| VVIn56 | 12 | 3.27 | 0.60 | 0.69 | 0.65 | 0.69 | 0.06 |
| VVIn52 | 13 | 7.87 | 0.58 | 0.87 | 0.86 | 0.87 | 0.20 |
| VVIq57 | 10 | 2.39 | 0.56 | 0.58 | 0.52 | 0.58 | 0.00 |
| VVIp31 | 24 | 11.14 | 0.88 | 0.91 | 0.90 | 0.91 | 0.01 |
| VVIp77 | 23 | 8.22 | 0.76 | 0.87 | 0.86 | 0.88 | 0.06 |
| VVIv36 | 15 | 4.74 | 0.79 | 0.78 | 0.76 | 0.79 | 0.00 |
| VVIr21 | 17 | 6.74 | 0.83 | 0.85 | 0.83 | 0.85 | 0.01 |
| VVS2 | 20 | 8.25 | 0.86 | 0.87 | 0.86 | 0.88 | 0.00 |
| VVMD5 | 18 | 8.80 | 0.76 | 0.88 | 0.87 | 0.88 | 0.07 |
| VVMD7 | 17 | 9.18 | 0.87 | 0.89 | 0.88 | 0.89 | 0.00 |
| VVMD25 | 19 | 5.91 | 0.75 | 0.83 | 0.81 | 0.83 | 0.04 |
| VVMD27 | 22 | 7.99 | 0.87 | 0.87 | 0.86 | 0.87 | 0.00 |
| VrZAG62 | 18 | 8.32 | 0.84 | 0.88 | 0.86 | 0.88 | 0.01 |
| VrZAG79 | 16 | 8.74 | 0.85 | 0.88 | 0.87 | 0.88 | 0.01 |
| **Total** | 304 | 119.42 |  |  |  |  |  |
| **Mean** | 17.88 | 7.02 | 0.75 | 0.83 | 0.81 | 0.83 |  |
| **SE\*** | 0.93 | 0.57 | 0.03 | 0.02 | 0.02 | 0.02 |  |

Number of alleles ( $N_a$ ), number of effective alleles ( $N_e$ ), observed heterozygosity ( $H_o$ ), expected heterozygosity ( $H_e$ ), polymorphic information content (PIC), discrimination power ( $D_j$ ), estimated frequency of null alleles ( $r$ ).
\*Standard error of mean values.

Across all the accessions, the mean observed heterozygosity (H_O_) was 0.75 (ranging from 0.48 to 0.88). The expected heterozygosity (H_E_) was higher than the observed heterozygosity (H_O_) for most loci, except for VVIv36. Among these loci (H_O_<H_E_), the probability of null alleles (*r*) was significantly high (>0.20) only for VVIp25b and VVIn52. The analysis revealed a high H_E_ level, ranging from 0.58 (VVIq57) to 0.91 (VVIp31), with a mean of 0.83.

The PIC estimates varied from 0.52 (VVIq57) to 0.90 (VVIp31), with a mean value of 0.81. The discrimination power (D*j*) was greater than 0.80 for 13 of the 17 loci, with the highest value for the VVIp31 locus (0.91). The D*j* values were high for 76.5% of the SSR markers used (>0.80). When the PIC and D*j* of each locus were analyzed together, 12 loci presented the highest values for both indexes (>0.80). In this study, the largest amount of information was provided by VVIp31, for which 24 alleles were detected showing a PIC and a D*j* ≥ 0.90.

### Evaluation of genetic relationships and germplasm structure

The STRUCTURE analysis indicated the relatedness among the 410 accessions, with the highest ΔK value for K = 3, suggesting that three genetic clusters were sufficient to interpret our data (Fig 1).

**Fig 1.**
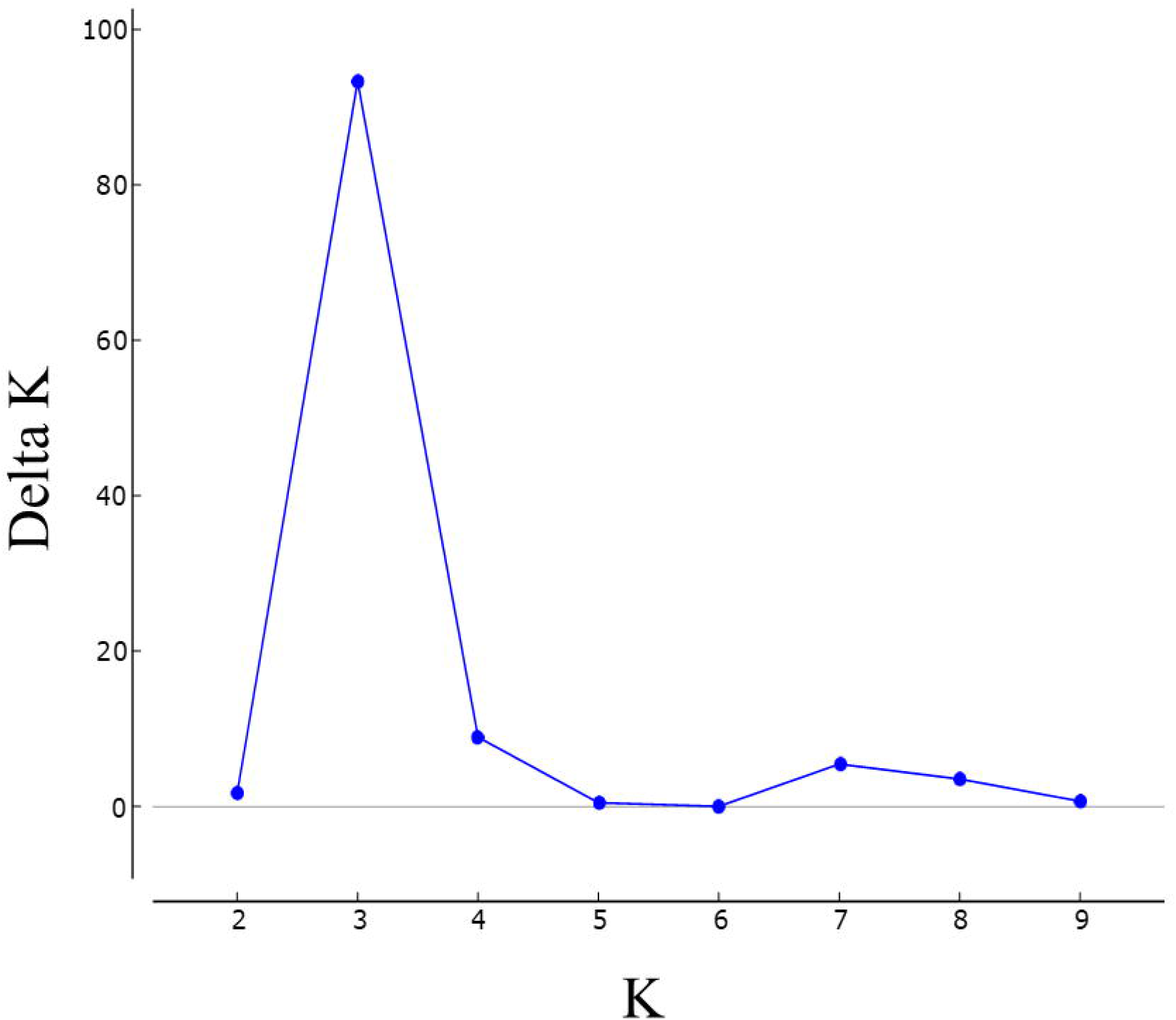
STRUCTURE Harvester results. The most probable number of genetic clusters (K) within the full data set of 410 individuals based on the method described by Evanno et al. (2005) [51]. Delta K graph determined the maximum value at K = 3.

Based on a membership probability threshold of 0.70, 207 accessions were assigned to cluster 1, 54 accessions were assigned to cluster 2, and 51 accessions were assigned to cluster 3. The remaining 98 accessions were assigned to the admixed group.

The level of clustering (K = 3) is related to the main accession species. Cluster 1 was formed by accessions with the greatest relation to *V. vinifera*. Cluster 2 contained the accessions most related to *V. labrusca*. Accessions linked to wild *Vitis* species were allocated to cluster 3. All accessions assigned to the admixed group were identified as interspecific hybrids (Fig 2A).

**Fig 2.**
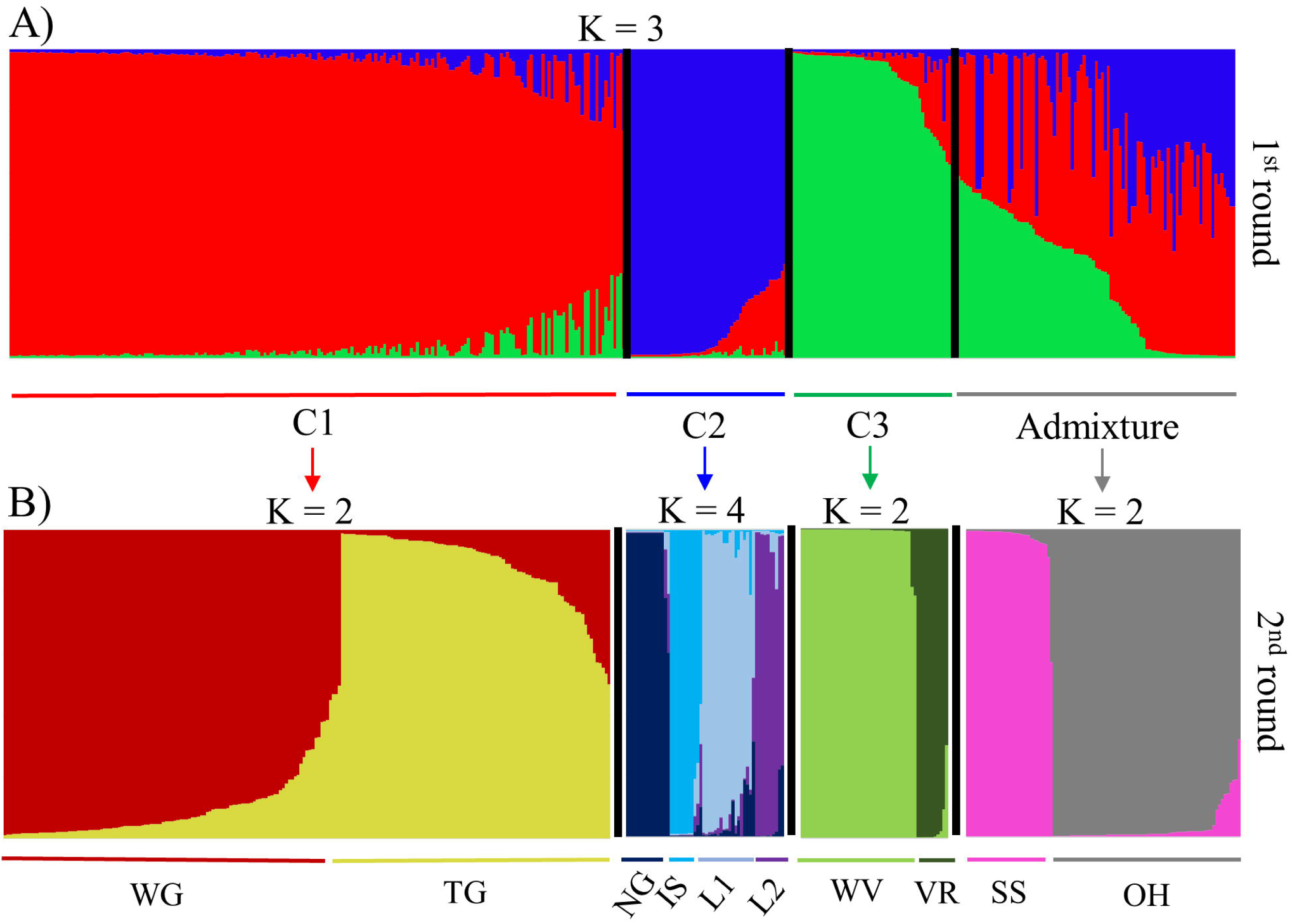
Genetic structure of the *Vitis* germplasm accessions obtained on the basis of 17 microsatellite markers. Bar graphs of the estimated membership proportions (q) for each of the 410 accessions. Each accession is represented by a single vertical line, which is partitioned into colored segments in proportion to the estimated membership in each cluster. [A] First round of STRUCTURE analysis, inferred genetic structure for K = 3. Cluster 1 (C1): genetic predominance of the species *V. vinifera*; cluster 2 (C2): genetic predominance of the species *V. labrusca*; cluster 3 (C3): genetic predominance of wild *Vitis* species; Admixture: interspecific hybrids with a membership of q < 0.70. [B] Second round of STRUCTURE analysis. WG: wine grape accessions related to *V. vinifera*; TG: table grape accessions related to *V. vinifera*; NG: ‘Niagara’ accessions; IS: ‘Ives’ and ‘Isabella’ accessions; L1 and L2: Others *V. labrusca* hybrids; WV: accessions related to wild *Vitis* species; VR: *V. rotundifolia* accessions; SS: accessions related to the Seibel series; OH: complex interspecific hybrids.

Of the 304 observed alleles, 227 were shared among the groups; the remaining 77 represented private alleles (Pa) in different groups of accessions (Fig 3). The VVMD27 locus had the largest number of private alleles of the 17 SSR markers used in this study (9). Clusters 1 and 2, constituted by accessions related to the most cultivated species of grapevine, *V. vinifera* and *V. labrusca,* respectively, had the smallest number of private alleles (5 and 1, respectively). The largest number of private alleles was found in cluster 3 (53), constituting 72.60% of the total private alleles.

**Fig 3.**
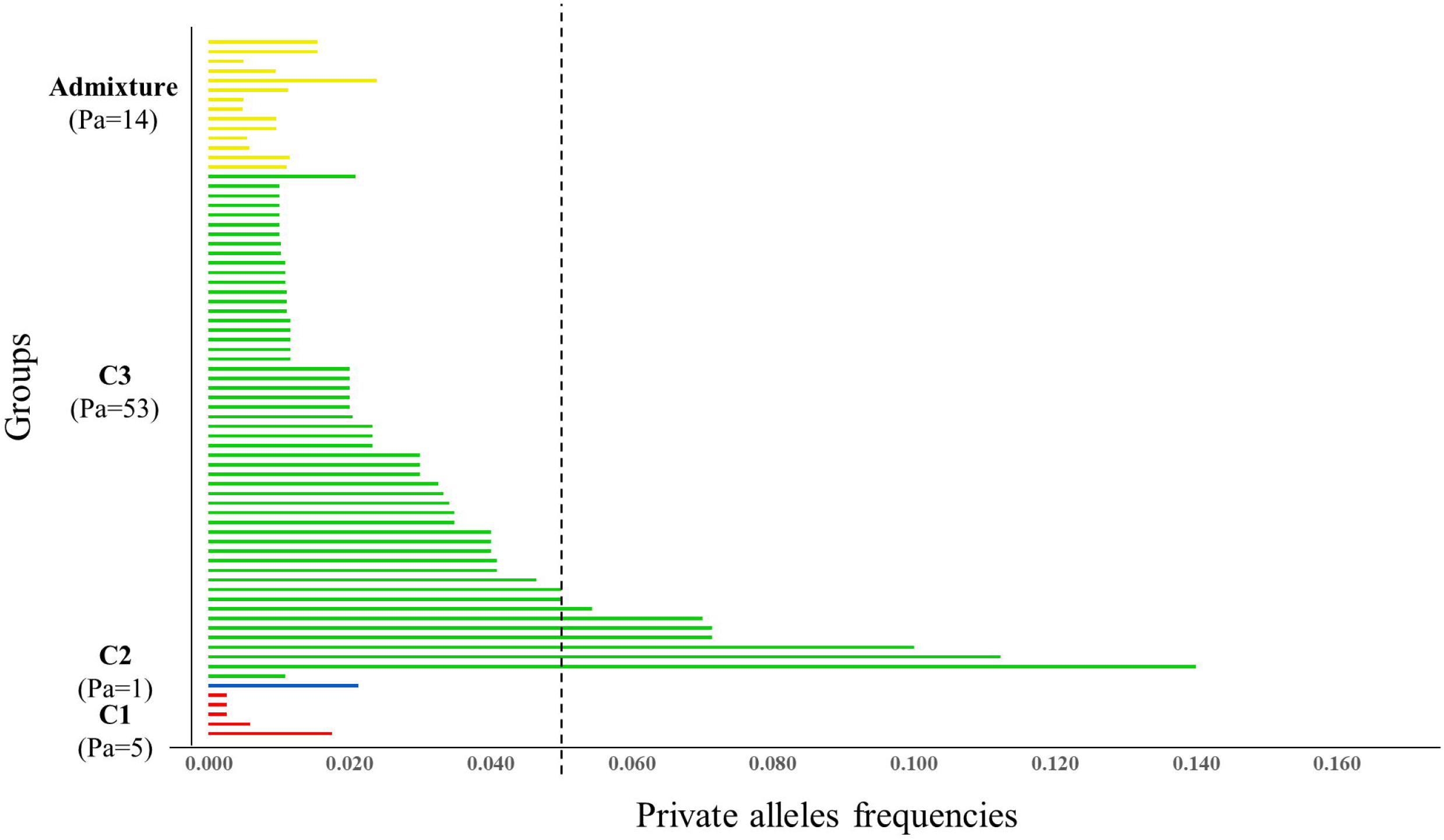
Private allele (Pa) frequencies obtained from the genotyping of 410 grapevine accessions on the basis of 17 microsatellite loci. X-axis: Private alleles frequencies; Y-axis: groups identified by STRUCTURE analyses at K = 3. The dashed line indicates the cutoff for the occurrence of rare alleles (frequency = 0.05).

A subsequent round (second round) of STRUCTURE allowed the identification of secondary clusters within the three main genetic clusters (Fig 2B). In Cluster 1, the accessions were divided into two subgroups (K = 2), one formed mainly by wine grapes (WG) (n = 115) and the other by table grapes (VT) (n = 92). This finer-scale clustering divided Cluster 2 into 4 subgroups (K = 4). The NG subgroup (n = 15) was composed of ‘Niagara’ and its mutations. In the IS subgroup (n = 11), the cultivars Ives, Isabella, and Isabella mutations were found. The remaining *V. labrusca* hybrids were allocated to subgroups L1 (n = 18) and L2 (n = 10). In cluster 3, the second round also divided the accessions into two subgroups (K = 2), the *V. rotundifolia* accessions were assigned to the VR subgroup (n = 11), and the others accessions related to wild *Vitis* species were allocated to the WV subgroup (n = 40).

Although the Admixture group contained a large number of heterogeneous accessions, a subsequent round of STRUCTURE was also performed on this set to identify possible clustering patterns. As a result, the analysis revealed the presence of two subgroups (K = 2). Accessions of the Seibel series and hybrids including cultivars of this complex in their genealogy were separated from the other hybrids and assigned to the SS subgroup (n = 31). The remaining 67 accessions of the Admixture group were in the OH subgroup.

Additionally, DAPC was performed with no prior information about the groupings of the evaluated accessions. Inspection of the BIC values (S1 Fig.) revealed that the division of the accessions into nine clusters was the most likely scheme to explain the variance in this set of accessions. In the preliminary step of data transformation, the maintenance of 120 principal components (PCs) allowed the DAPC to explain 94% of the total genetic variation.

Initially, the DAPC scatterplot based on the first and second discriminant functions showed the formation of three main distinct groups, with great genetic differentiation of clusters 8 (dark green) and 9 (green) from the others (Fig 4A). In a subsequent DAPC, outlier clusters 8 and 9 were removed to improve the visualization of the relationship of the other clusters (Fig 4B). In this second scatterplot, clusters 1 (magenta) and 7 (purple) showed greater genetic differentiation, with low variance within the groups, as well as no case of overlap with another cluster, indicating a strong genetic structure. The maintenance of 250 principal components (PCs) allowed the second DAPC to explain 100% of the total genetic variation.

**Fig 4.**
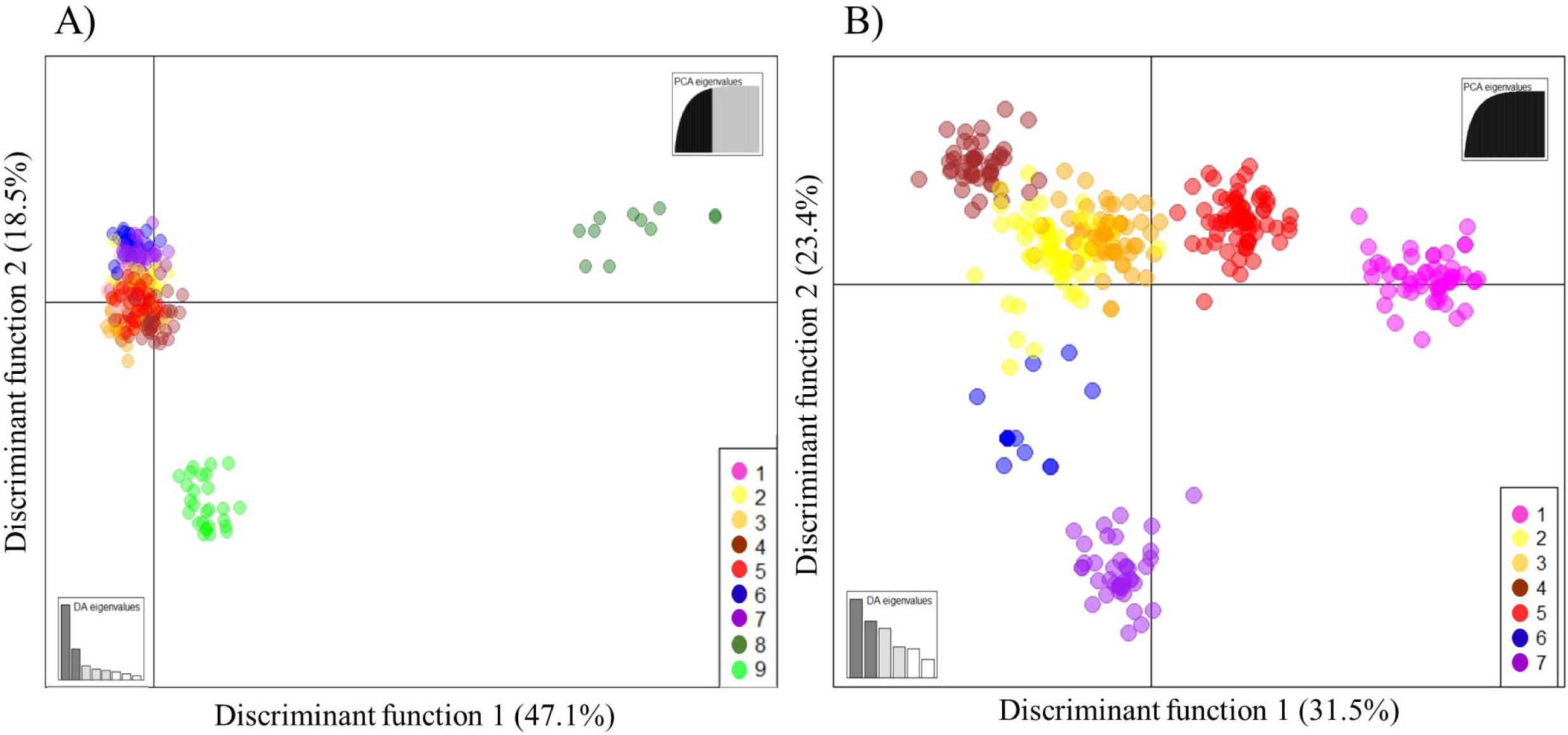
DAPC scatterplots based on the K-means algorithm used to identify the proper number of clusters. Dots represent individuals, and the clusters are presented in different colors. The accessions were allocated into nine clusters: 1 (magenta), related to the Seibel series; 2 (yellow), related to table grape accessions of *V. vinifera*; 3 (orange) and 5 (red), related to wine grape accessions of *V. vinifera*; 4 (brown), predominance of IAC hybrids; 6 (blue) and 7 (purple), related to the species *V. labrusca*; 8 (dark green), related to wild *Vitis* species; and 9 (green), *V. rotundifolia* accessions. [A] DAPC with all samples included. [B] DAPC excluding clusters 8 and 9.

The allocation of individuals into clusters according to the DAPC showed several similarities to those achieved in the second round of STRUCTURE, and both analyses showed the same pattern of clustering. Essentially, clusters 1 (magenta), 2 (yellow), 8 (dark green), and 9 (green) of the DAPC reflected the subgroups SS, TG, VR, and WV detected by the STRUCTURE second round, respectively, and the WG subgroup corresponded to DAPC clusters 5 (red) and 3 (orange).

In the case of the *V. labrusca* hybrids, the analyses resulted in a slightly different division. DAPC separated these accessions in clusters 6 (blue) and 7 (purple), basically assigning ‘Niagara’ accessions in cluster 6 and the other *V. labrusca* hybrids in cluster 7. The STRUCTURE second round also identified ‘Niagara’ accessions as a separate group (NG); however, a more refined division was performed in the other hybrids, separating them into 3 subgroups. DAPC cluster 4 (brown) did not correspond to any subgroup identified by the STRUCTURE second round; this cluster was formed mostly by hybrids developed by the IAC breeding program used as table grapes.

Finally, we constructed a dendrogram using the neighbor-joining method from the distance matrix based on Rogers’ distance to confirm the relationships among the accessions (Fig 5). The dendrogram showed a pattern that was consistent with those from the above-described two analyses. The group formed by the *V. rotundifolia* accessions and the other wild species was clearly separated from the cultivated *Vitis* species, as seen in the DAPC. There was also a strong separation between accessions related to *V. labrusca* and other accessions. The wine grape accessions of *V. vinifera* were mainly concentrated at the top of the dendrogram, while the table grape accessions of this species were found at the bottom. However, the other hybrids (IAC, Seibel series, and others) were scattered among all the groups formed by the dendrogram.

**Fig 5.**
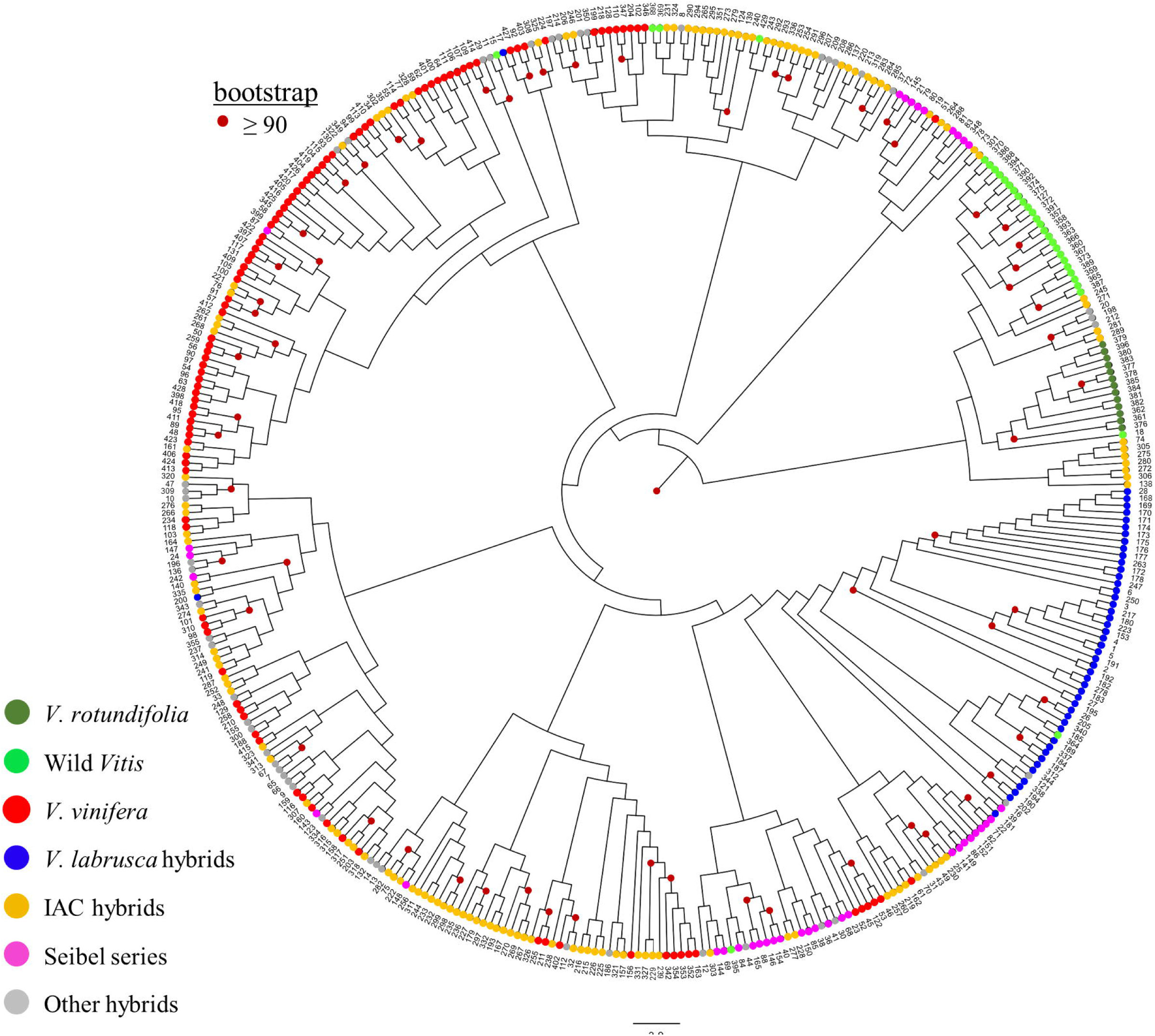
Neighbor-joining dendrogram based on Rogers’ distance calculated from the dataset of 17 microsatellite markers across 410 grapevine accessions. Accessions colored according to species group.

### Validation analysis of molecular profiles

The identification of 145 accessions was validated through matches with data available in the literature and databases. The results also confirmed matches to reference profiles of clones based on somatic mutations. Another 42 accessions showed molecular profiles that matched a validated reference profile of a different prime name, indicating mislabeling (S1 Table).

The molecular profiles of the remaining 223 accessions did not match any available reference profile. This accession group included wild species and cultivars from grapevine breeding programs in Brazil (the IAC and Embrapa), the United States, and France (Seibel series). The molecular profiles of more than 100 hybrids developed by the IAC were reported for the first time.

The accessions ‘101-14’, ‘Bailey’, ‘Black July’, ‘Carlos’, ‘Carman’, ‘Castelão’, ‘Catawba Rosa’, ‘Elvira’, ‘Moscatel de Alexandria’, and ‘Regent’ showed a different profile than the reference profile of the same name and did not match any other available reference profile. However, additional morphological and source information is needed to validate their identification. To avoid possible confusion, these accessions were indicated as “Unknown”.

After correcting the mislabeling, 22 cases of duplicates were identified, all with accessions of the same name and the same molecular profile. Accessions identified with different names but having the same molecular profile were classified as synonyms. Thirty-one synonymous groups were elucidated in this study (S3 Table). Some accessions classified as “Unknown” showed genetic profiles identical to accessions that did not match any available reference profile; examples can be seen in synonymous groups 1, 2, 5, and 6 in S3 Table.

### Core collection

Three independent sampling proportions were constructed with a size ranging from 10 to 30% of the entire dataset to identify the smallest set of accessions that would be able to represent as much of the available genetic diversity as possible (Table 2). Core 3, composed of 120 accessions, managed to capture 100% of the 304 detected alleles, while the smallest sample (Core 1) managed to capture 243 alleles, approximately 20% less than the total number of alleles detected. The genetic diversity index values obtained for the samples were similar to or higher than those for the entire germplasm. The H_O_ values ranged from 0.64 (Core 1) to 0.70 (Core 3); the value for Core 3 was similar to that detected for all 410 accessions (0.75). The three samples showed H_E_ and Ne values higher than those observed for the entire dataset. The H_E_ values for all samples were 0.85, while Ne ranged from 134.53 (Core 2) to 137.69 (Core 3). The values of Ne and H_E_ are related to allele frequencies, and low values of allele frequencies generate even lower values when squared. With a reduction in the number of accessions (N), the low-frequency alleles (allele frequency between 0.05 to 0.25) and rare alleles (frequency less than 0.05) showed an increase in frequency, resulting in an increase in Ne and H_E_.

**Table 2.** SSR diversity within each core collection compared with that of the entire dataset (IAC collection).

| Sample Name | Size | N | Na | Ne | $H_o^*$ | $H_E^*$ | Total SSR diversity captured (%) |
| --- | --- | --- | --- | --- | --- | --- | --- |
| Core 1 | 0.1 | 41 | 243 | 136.22 | 0.64 (0.03) | 0.85 (0.01) | 79.93 |
| Core 2 | 0.2 | 82 | 275 | 134.53 | 0.69 (0.04) | 0.85 (0.02) | 90.46 |
| Core 3 | 0.3 | 120 | 304 | 137.69 | 0.70 (0.03) | 0.85 (0.01) | 100 |
| IAC collection | 1.0 | 410 | 304 | 119.42 | 0.75 (0.03) | 0.83 (0.02) | 100 |

Number of accessions ( $N$ ), number of alleles ( $N_a$ ), number of effective alleles ( $N_e$ ), observed heterozygosity ( $H_o$ ), expected heterozygosity ( $H_E$ )
\*Standard error in parentheses.

Core 3 sample was the only one that managed to capture 100% of the alleles, being the best option for use in breeding as a core collection. All clusters detected in the STRUCTURE analysis and DAPC are represented in Core 3. In particular, in the STRUCTURE analysis at K = 3, 49 accessions were in cluster 1, 12 were in cluster 2, 29 were in cluster 3, and 30 were in the admixture group, representing 41, 10, 24, and 25% of Core 3, respectively.

## Discussion

### Genetic diversity

The results of this study revealed high levels of genetic diversity among the evaluated accessions. The observed high genetic diversity was expected since the grape germplasm from the IAC includes varieties with very diverse origins, wild species, and different intra- and interspecific hybrids.

We detected a H_E_ of 0.83 across the entire accession set in the 17 evaluated loci (Table 1). This result is similar to those found in other Brazilian germplasm banks characterized by containing European and American cultivars and an abundance of interspecific hybrids [64, 65]. However, this value was higher than that in the Iranian [11] (0.72), Turkish (0.75) [66], and Spanish (0.71) [67] collections, which possessed only *V. vinifera* accessions.

The large number of alleles per locus identified (∼18) was likely due to the taxonomic amplitude of the germplasm since a relatively large number of low-frequency alleles were found in wild species accessions. Lamboy and Alpha (1998) [17], when analyzing the diversity of 110 accessions belonging to 21 species of *Vitis* and 4 hybrids, detected 24.4 alleles per locus, a greater quantity than that observed in this study, showing that taxonomically broader accessions contribute to a greater number of alleles.

Most loci had lower H_O_ values than those expected from the randomized union of gametes (H_E_), except for VVIv36. For these loci, the probability of null alleles was positive but significantly high (> 0.20) for only VVIp25b and VVIn52. This finding suggests that at these loci, some of the apparent homozygotes could be heterozygous, with one allele being visible and the other not. Such null alleles can occur when mutations prevent the linking of primers to the target region [68].

The high number of alleles obtained by the 17 SSR primer set positively impacted the PIC and discrimination power (D*j*). PIC is an indicator of a marker’s informative ability in genetic studies (segregation, population identification, and paternity control), and its value reflects the polymorphism of the marker in the population studied. According to the classification of Botstein et al. (1980) [43], all the loci used can be considered highly informative (PIC > 0.50). The high D*j* values demonstrate that the microsatellite markers used in this study can be considered very effective for grape cultivar discrimination and could be valid to distinguish other accessions that could be introduced into the collection.

### Structure and genetic relationship of accessions

The genetic structure was mostly impacted by two factors that are difficult to separate: clear discrimination based on species and human usage as wine, table, or rootstock grapes, as previously noted by Laucou et al. (2018) [69] and Emanuelli et al. (2013) [70]. A population structure analysis using the software STRUCTURE revealed the presence of three primary clusters in our set of accessions based on the species *V. vinifera*, *V. labrusca,* and wild *Vitis*. This first structural level is also evidenced in the DAPC analysis and neighbor-joining dendrogram, where it is possible to observe a clear distinction of the accessions associated with *V. labrusca* and wild species.

However, a large number of accessions were not assigned and remained in a large admixed group, evidencing the genetic complexity of the analyzed plant material. Many of these accessions are crossbreeds between native vine species found in North America such as *V. riparia* Michaux, *V. rupestris* Scheele Michx, and *V. labrusca* L., and a number *V. vinifera* L. cultivars from Europe. The intra- and interspecific crossings carried out during breeding cycles in search of novelties and hybrid vigor promote the miscegenation of grapevine cultivars, resulting in hybrids with a heterogeneous genetic composition.

The assignment of these hybrids to groups based on species is often difficult, as these individuals certainly carry alleles from different gene pools, being in an intermediate position and belonging simultaneously to more than one cluster. The accessions ‘Campos da Paz’ and ‘IAC 0457-11 Iracema’ are examples of this condition. ‘Campos da Paz’ is an interspecific hybrid resulting from the cross between the cultivated species *V. vinifera* and the wild species *V. ruprestris*. The mixture of two genomes was detected by STRUCTURE, which assigned a membership probability threshold of 0.55 and 0.45 to clusters 1 and 3 respectively, representing the genetic clusters of the two parental species. A similar situation was observed for the accession ‘IAC 0457-11 Iracema’ developed from the cross between the species *V. vinifera* and *V. labrusca*, represented by genetic groups 1 and 2, respectively. The hybrid presented an intermediate membership of 0.5 to the two groups. The other accessions from Admixture group exhibited a similar or even more complex origin than these examples, and some of them were derived from crosses between more than three species, having associations with the three clusters simultaneously.

Our results demonstrated the largest number of private alleles in cluster 3 composed of the wild germplasm (Fig 3). This finding confirms that wild accessions are important reservoirs of genetic variation, with the potential for incorporating new materials into breeding programs in response to the demand for the development of cultivars with different characteristics. Wild grape germplasm is a potential source of unique alleles and provides the breeder with a set of genetic resources that may be useful in the development of cultivars that are resistant to pests and diseases, tolerant to abiotic stresses, and even show enhanced productivity, which makes their conservation of paramount importance [71].

The second round of STRUCTURE (Fig 2B) identified similar DAPC clustering patterns (Fig 4), in which the genotypes from *V. vinifera* were separated according to their use. The WG subgroup was composed mainly of wine grapes, such as the accessions ‘Syrah’, ‘Merlot Noir’, ‘Chenin Blanc’, ‘Petit Verdot’, and ‘Cabernet Sauvignon’, which showed associations with a membership greater than 0.95, corresponding to DAPC clusters 5 (red) and 3 (orange). The *V. vinifera* accessions of table grapes as ‘Centennial Seedless’, ‘Aigezard’, ‘Moscatel de Hamburgo’, and ‘Italia’ and their mutations ‘Benitaka’, ‘Rubi’, and ‘Brazil’ were found in the TG subgroup. This subgroup corresponded to cluster 2 (yellow) in the DAPC. In the neighbor-joining dendrogram, the *V. vinifera* accessions were also completely separated in terms of use; the wine grapes were located at the top, and the table grapes were located at the bottom. This result showed that the strong artificial selection based on human usage with wine or table influenced the genetic structure within the cultivated compartment of grapevine, as previously identified in previous studies [69, 72].

In the DAPC and neighbor-joining dendrogram, two groups were differentiated to a greater extent than the others (Fig 4 and Fig 5), with these groups being formed mainly of wild grapes that are often used as rootstocks. This phenomenon likely occurred because few rootstocks used worldwide contain part of the *V. vinifera* genome [9], while practically all table and wine grape hybrids present in this germplasm contain a part of it. In DAPC analyses, the *V. rotundifolia* accessions constituted the most divergent group. This species is the only one in the germplasm belonging to the *Muscadinia* subgenus, which contains plants with 2n = 40 chromosomes, while the others belong to the *Euvitis* subgenus, with 2n = 38 chromosomes. A high genetic divergence between *V. rotundifolia* and the species in the *Euvitis* subgenus was also observed by Costa et al. (2017) [73] through the use of RAPD molecular markers and by Miller et al. (2013) [74] through SNPs. The species *V. rotundifolia* is resistant to several grapevine pests and diseases [75] and is an important source of genetic material in the development of cultivars and rootstocks adapted to the most diverse environmental conditions and with tolerance and/or resistance to biotic and abiotic factors.

The DAPC cluster 1 was formed by only accessions of the Seibel series and hybrids with varieties of this series in their genealogy. The Seibel series is in fact a generic term that refers to several hybrid grapes developed in France at the end of the 19th century by Albert Seibel from crosses between European *V. vinifera* varieties and wild American *Vitis* species to develop phylloxera-resistant cultivars with characteristics of fine European grapes [76]. As these hybrids are derived from crosses among three or more species, most of them were identified as Admixture in the first round of the STRUCTURE. A second round of STRUCTURE was carried out in the Admixture group to confirm the structure of these accessions as shown by DAPC. As a result, the Seibel series accessions were separated from the other hybrids to form a subgroup, confirming the existence of a distinct gene pool. The combinations of alleles of different *Vitis* species clearly created unique genetic pools, with many related accessions, since they were developed using the same breeding program, which explains the grouping and genetic distinction.

The *V. labrusca* hybrids formed distinct groups in the three analyses. In the DAPC, this accession group was subdivided into two clusters (6 and 7) indicating the presence of a secondary structure between them (Fig 4). Cluster 6 contained only table grape cultivars, including ‘Eumelan’, ‘Niabell’, ‘Highland’, ‘Niagara’, and their mutations, while grape cultivars for processing, including ‘Isabella’, ‘Ives’, and ‘Concord Precoce’ were included in cluster 7. In the STRUCTURE second round, these accessions had a more pronounced division (Fig 2B), and the cultivars Niagara and Isabella together with their mutations were assigned to subgroups NG and IS, respectively, while the other accessions were distributed between subgroups L1 and L2. This refined secondary structuring was probably due to the hierarchical STRUCTURE method, since the sensitivity of the program is increased when using a primary cluster in isolation that allows for more detailed subdivisions [49].

The IAC breeding program started in 1943 with the aim of obtaining varieties of wine grapes, table grapes, and rootstocks. The first introductions in the Germplasm Bank constituted *V. viniferas* cultivars and Seilbel series hybrids originating in France. Subsequently, wild species and *V. labrusca* hybrids from North America were introduced. Varieties developed around the world continued to be introduced into the IAC germplasm (Table S1) over time, which currently has a large number of accesses originating mainly from the United States, France, and Italy, which correspond to 19.51%, 18.04%, and 8.78% of the germplasm, respectively. In smaller quantities, varieties from Argentina, Germany, Armenia, Spain, Japan, Portugal, and other countries are also found.

Many of the *V. vinifera* cultivars of the IAC germplasm originating in France, Italy, and Spain are common among grapevine germplasms worldwide, and their use in other studies of genetic diversity has been reported [67,69,77–80]. The American and Brazilian hybrids present in the germplasm are more restricted to collections in North and South America, being rarely reported in European studies [10,65,68].

With the results of the first crosses in the IAC breeding program, the hybrids with outstanding characteristics started to be used as parents [81]. Since the beginning of the program, more than 2,000 crosses have been performed over 50 years, using more than 800 parents [82]. Currently the germplasm has 134 accessions developed from these crossings, corresponding to 32.70% of the entire germplasm. Most of these hybrids are exclusive to this germplasm, and the molecular profiles of 109 are described for the first time in this study.

The broad genetic base and different objectives of the IAC breeding program were responsible for the development of hybrids with a wide genetic diversity, as evidenced in the three analyses revealing IAC hybrids in practically all the clusters. In the dendrogram, the IAC hybrids were highlighted to facilitate this perception (Fig 5). Over time, there has been a decrease in the importance of the wine industry in the State of São Paulo, and the search for table grape varieties has become predominant [82]. Some of these table grape hybrids developed by the IAC formed cluster 4 of the DAPC. The clustering of these hybrids is similar to the case of the Seibel series accessions, where the combinations of alleles from different crossings were probably responsible for the creation of a unique gene pool.

The analyses grouped most of the hybrids with one of their parents; however, cases in which the hybrids were not grouped with any of the parents occurred. Hybrids originating from the same crosses were not always grouped with the same parent. For example, the hybrids ‘IAC 0871-41 Patrícia’ and ‘IAC 0871-13 A Dona’ both resulted from the same crossing between hybrids ‘IAC 0501-06 Soraya’ and ‘IAC 0544-14’ located in DAPC clusters 2 and 3, respectively, hybrid ‘IAC 0871-41 Patrícia’ was grouped with its parent ‘IAC 0501-06 Soraya’, while ‘IAC 0871-13 A Dona’ was grouped with ‘IAC 0544-14’. These findings are easily explained when we consider the genetic biology of the grapevine. In general, grapevine cultivars are highly heterozygous, and crossing between divergent parents results in a highly segregating progeny. In the same progeny, the hybrids are heterogeneous, and they can present characteristics similar to both parents, similar to only one parent, or even different from both parents [83].

Since many of the accessions were introduced from different parts of the world and some others have a complex pedigree, it can be difficult to determine their true relationship. In the absence of information on the genetic relationships among most genotypes, it is not possible to determine the most accurate method of grouping. Although the use of multivariate techniques in the recognition of genetic diversity imposes a certain degree of structure in the data, and it is important to use different grouping criteria and the correct structure resulting from most of them to ensure that the obtained result is not an artifact of the technique used. The use of more than one clustering method, due to differences in hierarchization, optimization, and ordering of groups allows the classification to be complemented according to the criteria utilized by each technique and prevents erroneous inferences from being adopted in the allocation of materials within a given subgroup of genotypes [84].

The STRUCTURE grouping method could be contested because human manipulation of cultivars (displacements, breeding, clonal propagation) can generate a deviation from Hardy-Weinberg equilibrium; however, in our study, STRUCTURE analysis provided a very consistent attribution of genotypes to clusters. The Admixture group reflects the crossing among genotypes of the three groups identified in the first round of STRUCTURE corresponding to breeding activities in search of novelties and hybrid vigor. Furthermore, this analysis provides important information regarding the genetic composition of the hybrids, providing information about the proportion of each species in their genome. The three primary genetic groups of STRUCTURE were easily distinguished in the other analyses; however, in the DAPC analysis, new levels of structure were revealed within these primary groups. The DAPC analysis also provided information about the genetic divergence between the clusters, allowing the identification of related ones.

The STRUCTURE second round was carried out to investigate the presence of subclusters within the primary clusters and simultaneously validate the levels of structure obtained in the DAPC analysis. Most of the subgroups found in the STRUCTURE second round corresponded to the division obtained by the DAPC analysis, although some structural levels were different. These differences between analyses do not invalidate their results but rather bring complementary information that enhances understanding of the genetic structure and genetic relationship of germplasm accessions. The grouping based on the species of accessions was also evidenced by neighbor-joining dendrogram, but the differential of this analysis further provided visual information on the genetic relationship of the accessions within the groups. In the dendrogram, the genetic distance between two specific individuals was easily verified, providing a useful tool in breeding programs, mainly for the selection of divergent parents.

The information obtained by the STRUCTURE, DAPC, and neighbor-joining dendrogram provides important knowledge for the management of germplasm diversity. The identification of divergent groups guides crossings in breeding programs, facilitating the appropriate combination of parents to obtain progeny with wide genetic variability, allowing the maximization of heterosis and making it possible to obtain individuals with superior characteristics. Information about the available genetic diversity is valuable because if properly explored, it can reduce vulnerability to genetic erosion through the avoidance of crosses between genetically related genotypes while also accelerating genetic progress related to characteristics of importance to grape growth [85].

### Identification analysis: misnamed and synonymous cases

Considering the vast diversity of names for the different varieties of grapevine, standardization is necessary. Errors due to homonyms, synonyms, differences in spelling, and misnamed accessions impede estimation of the real number of different accessions that are present in grapevine collections, with a negative impact on grapevine breeding programs. Therefore, the verification of true-to-type accessions is indispensable [33]. For grapevine, a 7-SSR genotyping system has been established as a useful tool for identification and parentage analysis, allowing the allele length of varieties to be comparably scored by different institutions [34, 62].

In this study, 42 cases of misnaming were found by comparing the molecular profiles obtained with the information available in the literature and databases (S1 Table). The molecular profile of the accession ‘Cabernet Franc’ corresponded to the cultivar Merlot Noir, and the molecular profile of the accession ‘Merlot Noir’ corresponded to the cultivar Cabernet Franc, clearly indicating an exchange of nomenclature between these accessions. ‘Cabernet Franc’ is one of the parents of ‘Merlot Noir’, and some morphological traits of these two cultivars are quite similar [86], which certainly contributed to the occurrence of this mistake.

The molecular profile of the accession ‘Magoon’ matched that of ‘Regale’ in the present study, and ‘Regale’ had a similar molecular profile to those obtained by Schuck et al. (2011) [87] and Riaz et al. (2008) [88], indicating that ‘Magoon’ was misnamed at the time of introduction and that both accessions were the cultivar Regale. In Brazil, the same case of misnaming was also reported by Schuck et al. (2011) [87]. A misspelling case was observed for the accession ‘Pedro Ximenez’, corresponding to ‘Pedro Gimenez’; both cultivars are classified by the VIVC as wine grapes with white berries. However, despite the similar names and some comparable characteristics, the genealogies of these cultivars are completely different, being easily distinguished by microsatellite marker analysis due to the different molecular profiles generated.

The accessions ‘Armenia I70060’ and ‘Armenia I70061’ were labeled according to their country of origin, Armenia, during their introduction. Through microsatellite marker analysis, these accessions were identified as ‘Aigezard’ and ‘Parvana’, respectively. A similar situation was observed for the accession ‘Moscatel Suiça’, corresponding to ‘Muscat Bleu’ from Switzerland; this accession was likely also labeled according to its country of origin.

Additionally, 31 synonymous groups were identified (S3 Table). Cases of synonymy could correspond to clones of the same cultivar that show phenotypic differences due to the occurrence of somatic mutations [89, 90]. Mobile elements are known to generate somatic variation in vegetatively propagated plants such as grapevines [91, 92]. Carrier et al. (2012) [91] observed that insertion polymorphism caused by mobile elements is the major cause of mutational events related to clonal variation. In grape, retrotransposon-induced insertion into *VvmybA1*, a homolog of *VlmybA1-1*, is the molecular basis of the loss of pigmentation in a white grape cultivar of *V. vinifera* due to the lack of anthocyanin production [93].

The detection of somatic mutations is very difficult with a small number of microsatellite markers, especially when they are located in noncoding regions of the genome [94]. This was the case for synonymous groups 19, 20, 21, 22, in S3 Table, such as the cultivar Italia and its mutations ‘Rubi’, ‘Benitaka’, and ‘Brasil’, which differ in terms of the color of berries, with white, pink, red, and black fruits, respectively, and are cultivated as distinct cultivars in Brazil. This was also the case for ‘Pinot Gris’, a variant with gray berries arising from ‘Pinot Noir’, which has black berries. The mutations that occurred in the cultivar Niagara can be distinguished in terms of the color, size, and shape of the berries, and they may even lead to a lack of seeds, such as the apyrenic accession ‘Niagara Seedless’ or ‘Rosinha’ [95].

The accession ‘Tinta Roriz’ was identified as a synonym of the cultivar Tempranillo Tinto in this study; this synonym is already registered in the VIVC and is widely used in regions of Portugal [96]. The wild species *V. doaniana* and *V. berlandieri* have the same molecular profile, indicating a case of mislabeling; certainly, some mistake was made during the acquisition of these materials, and the same genotype was propagated with different names.

The occurrence of misidentification is common, especially for old clonal species such as *Vitis* spp., and it can occur during any stage of accession introduction and maintenance. It has been observed that 5 to 10% of the grape cultivars maintained in grape collections are incorrectly identified [97, 98]. In a new place, a certain genotype may receive a new name, confusing samples and the maintenance of accessions in germplasm banks [99]. The correct identification of accessions is fundamental to optimize germplasm management and for the use of germplasm in ongoing breeding programs since related genotypes will not be chosen for field experiments or controlled crosses. The identification of the existence of synonyms, homonyms, and misnamed accessions is essential to prevent future propagation and breeding errors [87] and in helping to reduce germplasm maintenance costs without the risk of losing valuable genetic resources. Since morphological descriptors are highly influenced by environmental factors, molecular analyses can support identification. SSR markers have often been considered very efficient at the cultivar level since they can be easily used to distinguish different cultivars; however, they are less effective in differentiating clones [9]. The results of molecular analysis should not replace ampelographic observations but should be integrated with such observations, mainly for the identification of somatic mutations.

In this study, 223 accessions with molecular profiles did not match any available reference profile. The largest subset of accessions was from the Brazilian grapevine breeding program of the IAC, with 109 molecular profiles described for the first time.

The identification and description of unreported molecular profiles is important for regional and national viticulture and ensures the institution’s intellectual property rights over these cultivars. The information obtained in this study will contribute to international cooperation to correctly identify grape germplasm and will allow the inclusion of new molecular profiles of Brazilian grapevine cultivars in the database.

### Development of a core collection

The intention for the development of a core collection is to represent the genetic diversity of the entire germplasm in a reduced set of accessions that is feasible to handle. The efficiency of the approach based on SSR profiles in identifying a core collection was already demonstrated for grapevine by Le Cunff et al. (2008) [100], Cipriani et al. (2010) [101], Emanuelli et al. (2013) [70] and Migliaro et al. (2019) [29].

In this study, 120 accessions (Core 3) were necessary to capture all the allelic diversity of the whole collection, which is equivalent to approximately 30% of all accessions (Table 2). In *V. vinifera* subsp*. sativa* core collections, the same result was obtained with smaller percentages of individuals, from 4 to 15% [70,100,101]. According to Le Cunff et al. (2008) [100], the use of only cultivated genotypes of *V. vinifera* subsp. *sativa* is one of the reasons for the small number of individuals in the core collection since cultivated genotypes tend to be less diverse than wild counterparts [12, 102].

Migliaro et al. (2019) [29] analyzed 379 grapevine rootstock accessions and managed to represent their full allelic richness with a core collection containing 30% of the accessions, a result similar to that observed in this study. According to these authors, the large number of individuals in the core collection can be related to the number of varieties belonging to different *Vitis* species and the high genetic variability detected. These are likely the same reasons for the need for a high number of genotypes in our core collection, since the *Vitis* spp. Germplasm Bank of the IAC also includes accessions belonging to different *Vitis* species and many interspecific hybrids that have complex pedigrees (derived from crosses among three or more species). The comparison of different methods used to form core collections is not easy, as the analyses are rarely performed in the same way, and the original collections rarely include the same global diversity of species [100].

Among the 120 genotypes in Core 3, 82 were identified as interspecific hybrids, with 13 being non-*vinifera* varieties. This large number of interspecific hybrids in the core collection can be explained by their predominance in germplasm; in addition, many of them have a complex pedigree, which certainly combines alleles of different species of *Vitis*. Regarding the other genotypes in Core 3, 31 were identified as *V. vinifera* cultivars and seven as wild *Vitis*.

The core collection was constructed to provide a logical subset of germplasm for examination when the entire collection cannot be used. Complementary criteria, such as phenotypic, agronomic, and adaptive traits, should be associated with the core collection to make it more fully representative. Finally, this core collection will be useful for the development of new breeding strategies, future phenotyping efforts, and genome-wide association studies.

## Conclusions

A wide range of genetic diversity was revealed in the studied germplasm, ensuring the conservation of a large portion of grapevine genetic resources. The genetic diversity showed a pattern of structuring based on the species and use of accessions, as evidenced in a manner similar to the three structuring analyses. In addition, each of the analyses provided different information that was be complementary and equally valuable for breeding.

Taken together, our results can be used to efficiently guide future breeding efforts, whether through traditional hybridization or new breeding technologies. The obtained information may also enhance the management of grapevine germplasms and provide molecular data from a large set of genetic resources that contribute to expanding existing database information.

## Supporting information

S1 Table. Genotypes used in this study.

S2 Table. Name, linkage group, microsatellite sequences, and references of the SSR markers used in this study.

S3 Table. List of synonyms found in the Vitis spp. Germplasm Bank of the Agronomic Institute of Campinas (IAC) by SSR analysis.

S1 Fig. Bayesian information criterion (BIC) values for different numbers of clusters.

S2 Fig. Harvester results for STRUCTURE second round.

## Acknowledgments

The authors are grateful to the São Paulo Research Foundation (FAPESP) and Coordination for the Improvement of Higher Education Personnel (CAPES) for supporting the project and researchers.

## Supporting Information

**S1 Table. Genotypes used in this study.** Detailed characteristics: SSR matches, usage, country of origin, species, group assignments according to the DAPC, and STRUCTURE first and second round, core collection composition, and genotyping results obtained with the 17 microsatellite markers.

**S2 Table. Name, linkage group, microsatellite sequences, and references of the SSR markers used in this study.**

**S3 Table. List of synonyms found in the *Vitis* spp. Germplasm Bank of the Agronomic Institute of Campinas (IAC) by SSR analysis.**

**S1 Fig. Bayesian information criterion (BIC) values for different numbers of clusters.** The accepted true number of clusters was nine.

**S2 Fig. Harvester results for STRUCTURE second round.** Graphics for the detection of the most probable number of groups (K) estimated based on the method described by Evanno et al. (2005) [51]. [A] Cluster 1 - Highest peak for K = 2. [B] Cluster 2 - Highest peak for K = 4. [C] Cluster 3 - Highest peak for K = 2. [D] Admixture group - Highest peak for K = 2.

## Notes

### Competing Interest Statement

The authors have declared no competing interest.

