## Supplementary material for "Genetic structure and molecular diversity of Brazilian grapevine germplasm: management and use in breeding programs": S2 Table. Name, linkage group, microsatellite sequences, and references of the SSR markers used in this study.

| **SSR locus Name** | **Linkage Group**^a^ | **Microsatellite Repeat Motif** | **Reference** |
| --- | --- | --- | --- |
| VVIn74 | 19 | (AG)_11_ | Merdinoglu et al. (2005) [32] |
| VVIr09 | 18 | (GA)_19_ | Merdinoglu et al. (2005) [32] |
| VVIp25b | 4 | (GA)_14_ | Merdinoglu et al. (2005) [32] |
| VVIn56 | 7 | (AC)_9_ | Merdinoglu et al. (2005) [32] |
| VVIn52 | 16 | (GA)_11_ | Merdinoglu et al. (2005) [32] |
| VVIq57 | 1 | (GT)_6_ | Merdinoglu et al. (2005) [32] |
| VVIp31 | 19 | (GA)_20_ | Merdinoglu et al. (2005) [32] |
| VVIp77 | 4 | (CT)_11_ | Merdinoglu et al. (2005) [32] |
| VVIv36 | 7 | (GA)_10_ | Merdinoglu et al. (2005) [32] |
| VVIr21 | 10 | (GA)_11_ | Merdinoglu et al. (2005) [32] |
| VVS2 | 11 | (GA)_22_ | Thomas e Scott (1993) [35] |
| VVMD5 | 16 | (CT)_3_AT(CT)_11_ATAG(AT)_3_ | Bowers et al. (1996) [36] |
| VVMD7 | 7 | (CT)_14,5_ | Bowers et al. (1996) [36] |
| VVMD25 | 11 | (CT)_n_ | Bowers et al. (1999) [37] |
| VVMD27 | 5 | (CT)_n_ | Bowers et al. (1999) [37] |
| VrZAG62 | 7 | (GA)_19_ | Sefc et al. (1999) [38] |
| VrZAG79 | 5 | (GA)_19_ | Sefc et al. (1999) [38] |

^a^ Linkage groups are numbered according to Adam-Blondon et al. (2004) [103], Merdinoglu et al. (2005) [32] and Zarouri et al. (2015) [30].
