## Supplementary material for "Genetic structure and molecular diversity of Brazilian grapevine germplasm: management and use in breeding programs": S3 Table. List of synonyms found in the Vitis spp. Germplasm Bank of the Agronomic Institute of Campinas (IAC) by SSR analysis.

| **Group** | **Accession numbers** | **Synonyms** |
| --- | --- | --- |
| 1 | 310-98-101 | Armênia I70058, Unknown 06, Unknown 09 |
| 2 | 15-11 | Cunningham, Unknown 03 |
| 3 | 184-187 | Dona Zilá, Tardia de Caxias |
| 4 | 77-328 | SR 496-15, IAC 0496-15 |
| 5 | 205-26 | Fern Munson, Unknown 07 |
| 6 | 202-190 | Hidalgo, Unknown 05 |
| 7 | 251-317 | IAC 0506-33 (Isaura), IAC 1596-02 (Marta) |
| 8 | 253-254 | IAC 0544-14, IAC 0547-02 |
| 9 | 267-269 | IAC 0746-03, IAC 0740-01 |
| 10 | 206-246 | IAC 0775-26 (Aurora), IAC 0457-11 (Iracema) |
| 11 | 283-284 | IAC 0871-05 (Geni), IAC 0871-13 (A Dona) |
| 12 | 293-336 | IAC 0904-30, IAC 0733-39 |
| 13 | 298-299 | IAC 0915-02, IAC 0918-52 |
| 14 | 322-349 | IAC 1410-04, Romana |
| 15 | 124-139 | IAC 1848-04, IAC 1848-11 |
| 16 | 140-242 | IAC 1897-16, Seibel 07144 |
| 17 | 229-327 | IAC 387, IAC 486-03 |
| 18 | 261-262 | IAC 592-01, IAC 594-03 |
| 19 | 1-4-5-223 | Isabel, Isabelão, Isabel Precoce, Izabel Sport |
| 20 | 352-163-353-354 | Itália, Brasil, Rubi, Benitaka |
| 21 | 168-169-170-171-172-173-174-175-176-177-178 | Niagara Rosada, Niagara Branca, Niagara Rajada, Niagara sem sementes Rosinha, Niagara Rosada Gigante, Niagara Steck, Niagara Rosada Escura, Niagara Rosada Variegada, Niagara Branca Oval, Niagara Maravilha, Niagara Branca Gigante |
| 22 | 114-55 | Pinot Gris, Pinot Noir |
| 23 | 34-302 | Rainha, IAC 966-01 |
| 24 | 72-145 | Seibel 1077, Seibel 6086 |
| 25 | 40-154 | Seibel 13680, Seibel 13693 |
| 26 | 79-80 | Seibel 159, Seibel 848 |
| 27 | 88-165 | Seibel 5163, Seibel 5593 |
| 28 | 75-282 | SR 5010-08, SR 5010-21 |
| 29 | 25-230 | SR 5012-34 (Dona Emília), SR 507-08 |
| 30 | 104-115 | Tempranillo, Tinta Roriz |
| 31 | 363-366 | *Vitis doaniana*, *Vitis berlandieri* |
