## Supplementary figures and images for "Genetic structure and molecular diversity of Brazilian grapevine germplasm: management and use in breeding programs"

### S1 Fig. Bayesian information criterion (BIC) values for different numbers of clusters.

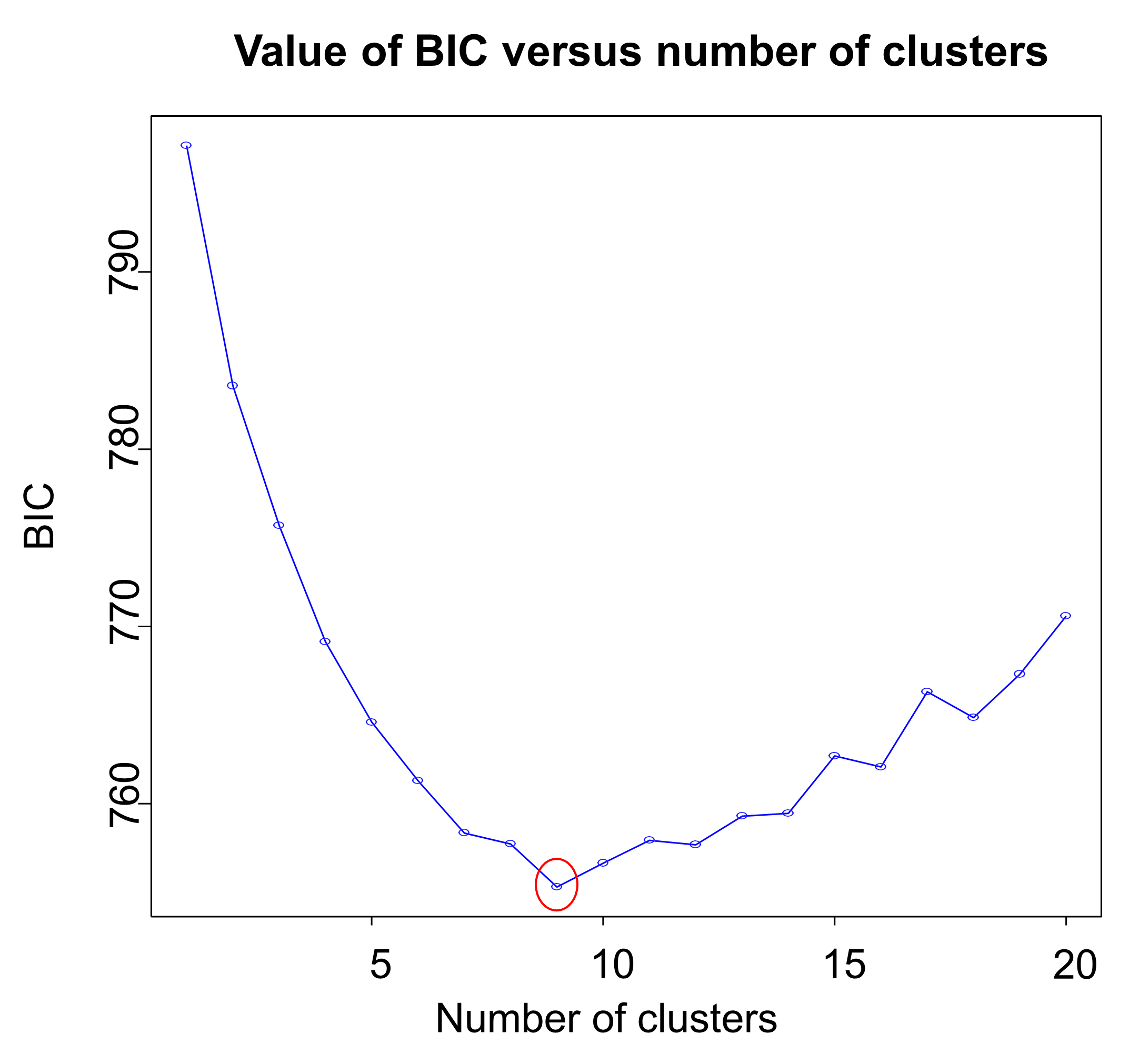

### S2 Fig. Harvester results for STRUCTURE second round.

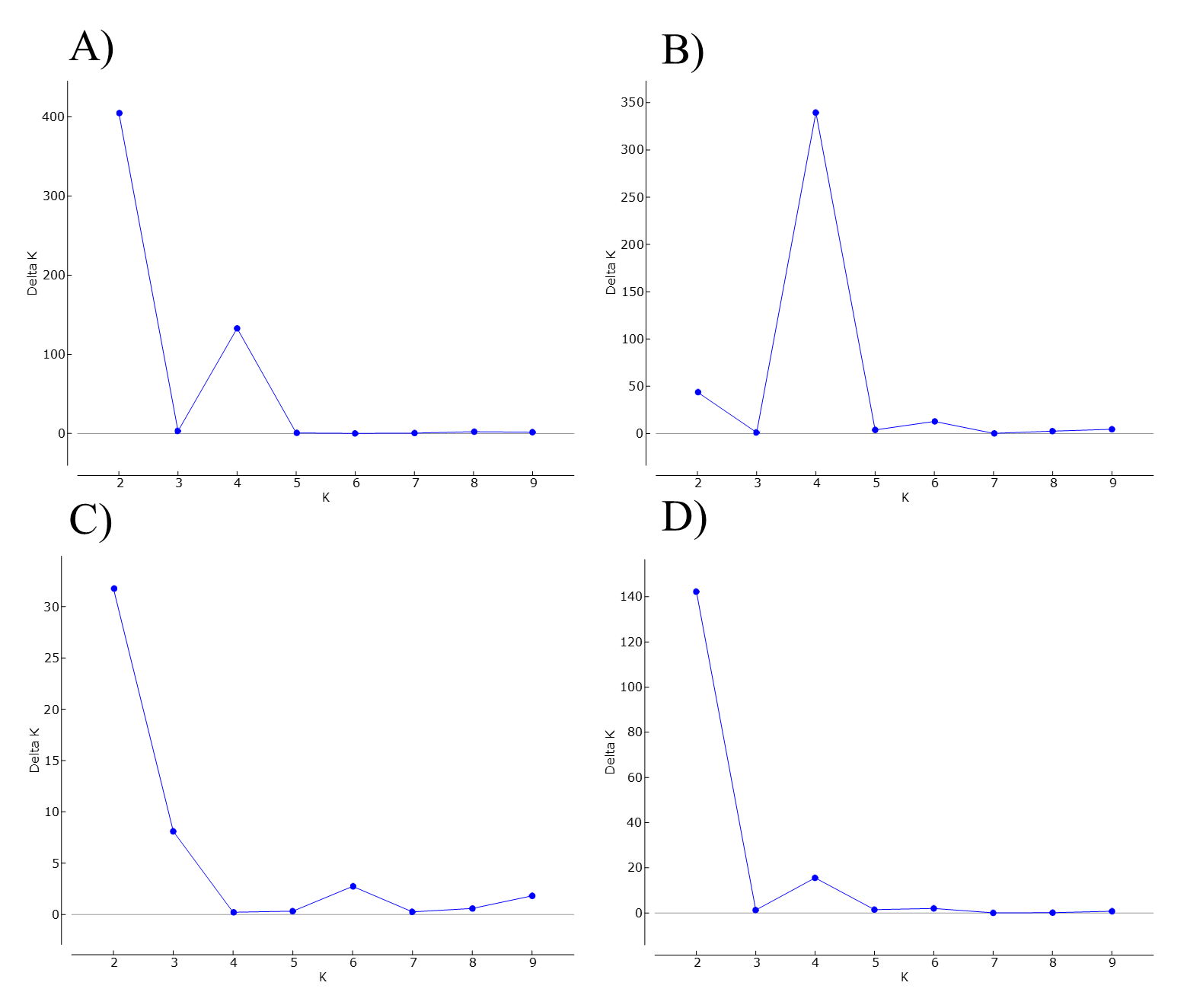
